# Brassinosteroid recruits FERONIA to safeguard cell expansion in Arabidopsis

**DOI:** 10.1101/2023.10.01.560400

**Authors:** Ajeet Chaudhary, Yu-Chun Hsiao, Fang-Ling Jessica Yeh, Hen-Ming Wu, Alice Y. Cheung, Shou-Ling Xu, Zhi-Yong Wang

## Abstract

Plant cell expansion is driven by turgor pressure and regulated by hormones. How plant cells avoid cell wall rupture during hormone-induced cell expansion remains a mystery. Here we show that brassinosteroid (BR), while stimulating cell elongation, promotes the plasma membrane (PM) accumulation of the receptor kinase FERONIA (FER), which monitors cell wall damage and in turn attenuates BR-induced cell elongation to prevent cell rupture. The GSK3-like kinase BIN2 phosphorylates FER, resulting in reduced FER accumulation and translocation from endoplasmic reticulum to PM. By inactivating BIN2, BR signaling promotes dephosphorylation and increases PM accumulation of FER, thereby enhancing the surveillance of cell wall integrity. Our study reveals a vital signaling circuit that coordinates hormone signaling with mechanical sensing to prevent cell bursting during hormone-induced cell expansion.

**One-Sentence Summary:** Brassinosteroid recruits a cell wall integrity monitor to prevent growth-induced cell wall damage.

## Main Text

The cell wall presents unique challenges for plant growth regulation. Cell expansion requires turgor pressure, weakening of connections between cell wall components, and synthesis of cell wall materials. Maximizing cell expansion requires precise regulation that balances these three processes; an imbalance would lead to either suboptimal growth or cell wall rupture by turgor pressure. Therefore, a crucial aspect of growth regulation is to coordinate between signalling pathways that respond to growth-promoting signals and those that monitor cell wall mechanics and integrity (*1*).

Brassinosteroid (BR) is a growth hormone that plays a major role in increasing plant size and biomass. While BR plays diverse cell-type specific roles in plant development (*2*), its main effect on plant size is from its promotion of epidermal cell expansion (*3*), which in turn allows expansion of internal tissue (*4*). BR binds to the receptor kinase BRI1, which activates a signaling cascade that leads to the inactivation of the GSK3-like kinase BIN2 through tyrosine dephosphorylation and ubiquitination-mediated degradation (*5-7*). Inactivation of BIN2 results in the dephosphorylation and activation of the BZR1 family transcription factors, which mediate transcriptional activation of growth-promoting genes including genes involved in cell wall synthesis and remodeling (*6*). BR also induces phosphorylation and activation of PM-H^+^-ATPase, which increase turgor pressure and cell wall acidity to promote cell expansion (*8, 9*). BR has also been implicated in plant responses to the imbalance in wall components by regulating cell wall biosynthesis (*1, 10*). A recent study reveals that the coordination between BR signaling and mechanical cues is essential for both morphogenesis and tissue integrity (*4*). However, the molecular mechanisms of such coordination are unknown.

Receptor kinases that sense cell wall defects have been identified (1). However, their role in maintaining cell integrity during hormone-induced cell expansion and growth remains unknown. In this study, we tested whether wall-sensing receptors are required for maintaining cell wall integrity (CWI) during BR-induced cell expansion and growth. We uncovered not only an essential role for the FERONIA receptor kinase in preventing cell wall damage during BR-induced growth but also a signaling circuit by which BR recruits FER to enhance cell wall surveillance, potentially balancing different components of cell expansion to prevent cell bursting.

## FER is required to avoid cell bursts during BR-induced growth

To test the role wall-sensing receptors play in maintaining cell wall integrity during BR-induced cell expansion, we analyzed how various concentrations of BR affect the growth, morphology, and cell integrity of mutants defective in cell wall-sensing receptor kinases. These include FERONIA (*fer-4*) (*11*), HERCULES1(*herk1-1) (12*), THESEUS1 (*the1-1*), STRUBBELIG (*sub-21), and* MALE DISCOVERER1-INTERACTING RECEPTOR LIKE KINASE 2 (*mik2-1)(13*). We grew the mutants and wild-type (WT) seedlings on various concentrations (1, 10, 100, 1000 nM) of brassinolide (BL, the most active form of BR) (fig. S1). BL increased the hypocotyl elongation and reduced root growth of the WT in a dose-dependent manner. All the mutants except *fer-4* showed similar BR responses to WT (fig. S1). The roots of the *fer* mutants were of similar length as WT on medium without BL, slightly longer than WT on low concentrations of BL (1 and 10 nM), and shorter than WT on high concentrations of BL (100 and 1000 nM) (Fig. 1, A and B). The hypocotyls of *fer* mutants were longest on 100 nM BL and were longer than WT at 10 nM and 100 nM of BL but shorter at 1000 nM BL (Fig. 1, A and C). We also noticed that treatment with high concentrations of BL caused *fer* seedlings to accumulate increasing levels of anthocyanin pigment, which is usually caused by stress (Fig. 1, A and D). The *fer* mutant complemented by a *FER-GFP* transgene (*FER-GFP/fer*) showed the same phenotypes as the WT. These results suggest that *fer* is hypersensitive to BL and experiences stress on high concentrations of BL.

**Fig. 1.**
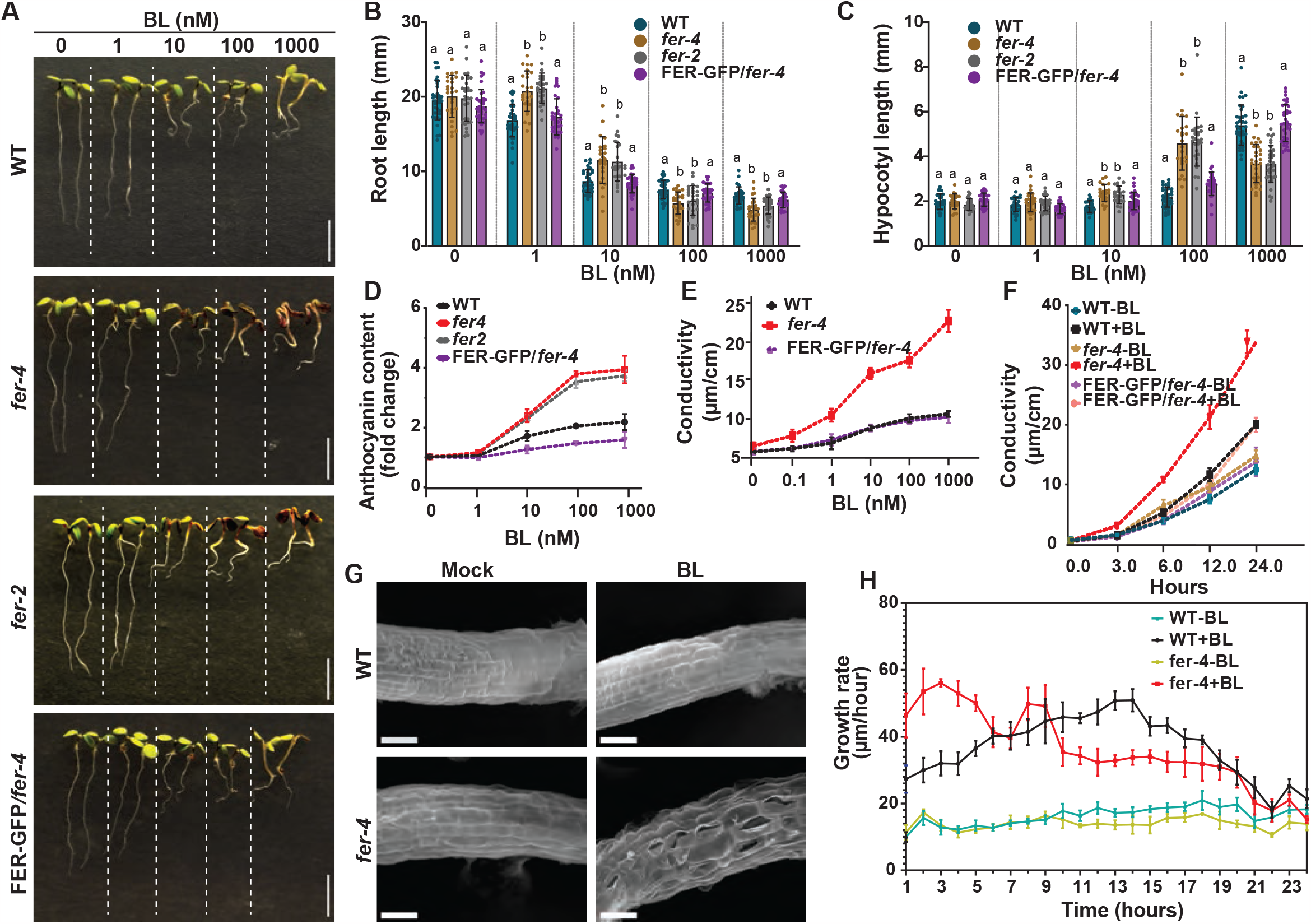
FER is essential for maintaining cell wall integrity during BR-induced cell elongation. (**A**) Images of seedlings of WT, *fer-2, fer-4* and *FER-GFP/fer-4* grown under 16 h 25 light/8 h dark cycle for 6 days on solid media containing the indicated concentrations of BL. Scale bars are 10 mm. (**B, C**) Quantification of root (B) and hypocotyl (C) lengths of seedlings shown in (A). Results presented here are mean ± SD of measurements pooled from three independent experiments (22≤N≤36 seedlings). Statistical significance was calculated using one-way ANOVA followed by Tukey’s test. Means with different letters are significantly different at P < 0.0001. (**D**) Anthocyanin accumulation in 6-day-old seedlings of WT, *fer-2, fer-4* and *FER-GFP*/*fer-4*. Data were normalized to the wild-type sample grown without BL (mean ± SD (N=3)). The experiment was repeated three times with similar results. (**E** and **F**) Electrolyte leakage in WT, *fer-4, FER-GFP*/*fer-4* plants treated with various concentrations of BL for 12 hours (E) or treated with 1 μM BL for various times (F). Values are the means ± SE of three biological replicates. The experiment was performed three times with similar results. (**G**) Scanning electron microscope (SEM) images of WT and *fer-4* primary roots treated with 1 μM BL or mock solution. Scale bars are 50 μm. (**H**) The root growth rates of WT and *fer-4* plants grown for 4 days without BL and then transferred to media containing no BL (-BL) or 1 μM BL (+BL). Values are mean ± SD (N=5). The experiment was performed two times with similar results.

We further tested whether overexpression of FER or its ligand RALF1 peptide had the opposite effects of loss of function *fer* mutation. In the absence of exogenous BL, the transgenic plants that overexpress FER (*FER-OE*, confirmed by RT-PCR, fig. S2A) and RALF1 (*RALF1-OE*) showed reduced root growth and marginally reduced hypocotyl growth compared to WT (fig. S2, B-D). At high concentrations (100 and 1000 nM) of BL, the *FER-OE* lines showed longer root and hypocotyl than WT and *fer-4*. The *ralf1* mutant and *RALF1-OE* showed similar growth responses to BL as WT. Moreover, the *FER-OE* and *RALF1-OE* lines showed reduced red pigmentation (anthocyanin) on BL (fig. S2E). Together the loss- and gain-of-function phenotypes consistently suggest that FER attenuates BR-induced growth and reduces stresses caused by BR.

To determine whether high concentrations of BL caused cell damage in *fer*, we measured electrolyte leakage of the plants grown on various concentrations of BL (Fig. 1E,). We grew seedlings on solid media for four days and then transferred the seedlings to deionized water supplemented with 1% sucrose and various concentrations of BL for 24 hours. Leakage of cellular content was measured as the electric conductivity of the water after seedling incubation relative to the total conductivity after boiling the seedlings (*14*). Increasing concentrations of BL or time of 1000 nM BL treatment caused much higher electrolyte leakage in the *fer* seedlings than in WT (Fig. 1E, F). To confirm the cell damage, we analyzed the WT and *fer* root tips using scanning electron microscopy (SEM). We observed extensive cell bursts in the elongation zone of the BR-treated *fer*, but not in wild-type or untreated *fer* mutant (Fig. 1G).

To understand whether FER prevents cell damage by slowing down cell elongation upon BR stimulation, we measured the growth kinetics after transferring seedlings from BR-free media to media containing 0 or 1000 nM BL (Fig. 1H). On media containing no BL, the WT and *fer* roots grew at a comparable rate of about 15 um/hour. Upon transferring to BL media, the growth rate of WT roots slowly increased from about 27 μm/hr to 50 μm/hr in about 14 hrs and then declined. This BR response kinetics is consistent with the previous report (*2*). By contrast, the roots of *fer-4* mutant responded to BL rapidly with a growth rate reaching 46 μm/hr within 1 hr, a maximum 55 μm/hr at 3 hrs of BL treatment and declined afterward. These results indicate that FER ensures a slow acceleration in cell elongation upon BR stimulation. The unabated rapid cell elongation likely results in cell wall damage and cell bursting in the BR-treated *fer*.

## BR promotes FER accumulation at the plasma membrane

FER is the most extensively studied member of the malectin domain-containing receptor-like kinase (MLD-RLK) (also known as Catharanthus roseus RLK1-LIKE or CrRLK1L) family. It plays broad and profound roles in diverse processes including sexual reproduction, development of root hair, hormonal responses, mechanic sensing, plant-microbe interactions, and maintaining CWI under abiotic stresses (*15-24*). Previous studies have shown that BR increases *FER* RNA level (*12*). However, we recently identified FER as a substrate of the BR-signaling GSK3 kinase BIN2 (*25*), suggesting that BR may regulate FER through a rapid posttranslational mechanism. To understand how BR signaling regulates FER, we treated *pFER:FER-GFP* seedlings with BL for various time periods and analyzed FER RNA and protein levels. The qRT-PCR analysis showed an increase of *FER* RNA after 1 hr but not 30 min of BR treatment (Fig. 2A). In contrast, immunoblot analysis showed that the FER protein level increased after 15 min of BR treatment, before the increase of its RNA level (Fig. 2B), suggesting posttranscriptional regulation. Treating the seedlings with bikinin, an inhibitor of BIN2 kinase, caused a similar increase of FER protein level (Fig. 2C), indicating that BIN2 mediates the BR regulation of FER protein accumulation. We repeated the experiment using transgenic plants that express FER-GFP from the constitutive 35S promoter (*35S:FER-GFP*), and the results confirmed that BR increases FER level posttranscriptionally (Fig. 2D).

**Fig. 2.**
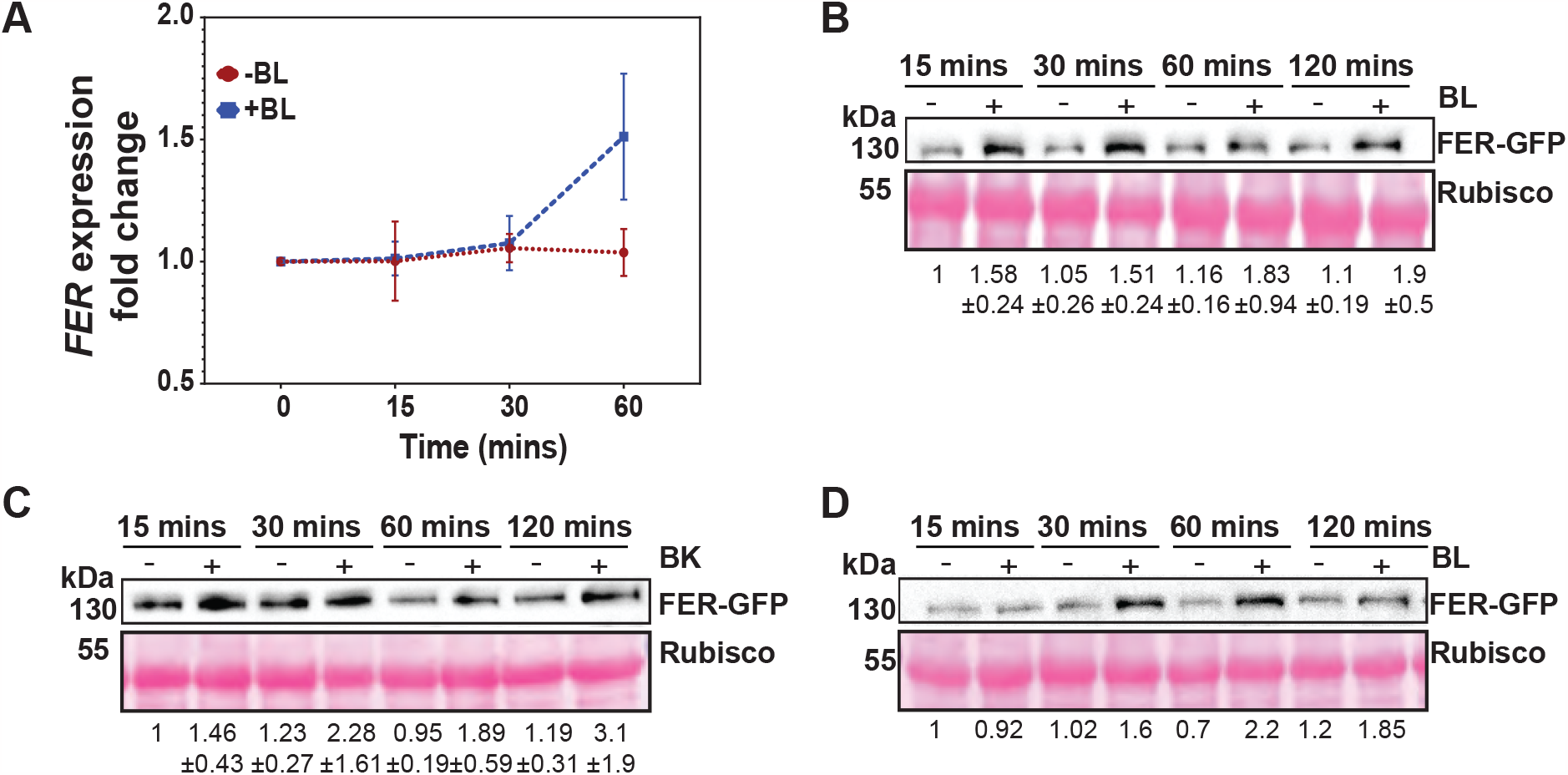
BL increases FER protein level through both transcriptional and post-transcriptional mechanisms. (**A**) RT-qPCR analysis of *FER* expression after BL treatment. Values are the means ± SD of three biological replicates. (**B-D**) Immunoblot analysis of FER-GFP protein in *pFER:FER-GFP/fer-4* (B, C) or *35S:FER-GFP* (D) plants after various treatment times with 1000 nM BL (B and D) or 50 μM bikinin (C). The top panel shows anti-GFP immunoblots and the bottom panel is ponceau-stained rubisco as a loading control. Numbers below the images are quantification (means ± SD) of FER-GFP signals from three independent replicates.

Next, we analyzed BR’s effects on sub-cellular localization and distribution of FER. Our confocal microcopy analysis showed that FER-GFP is localized to the plasma membrane and endoplasmic reticulum (Fig. 3A), consistent with previous reports (*11, 26*). Treating the seedlings with 1 μM propiconazole (PPZ, an inhibitor of BR biosynthesis) for 24 hrs reduced FER-GFP signal at the plasma membrane and enhanced cytoplasmic signal of FER-GFP colocalizing with ER-RFP (Mander’s overlap coefficient 0.58). BL treatment increased FER-GFP at the plasma membrane and reduced the cytoplasmic FER-GFP that colocalized with ER-RFP (Mander’s overlap coefficient less than 0.2). Bikinin, the BIN2 inhibitor, had a similar effect as BL (Fig. 3A, B). We found that within 30 min of BL or bikinin treatment, the colocalization of FER-GFP with ER-RFP was greatly reduced (Fig. 3C and D). Moreover, co-expression of BIN2 in protoplast caused ER retention of FER-GFP in a dose-dependent manner and the effect of BIN2 was reversed by BR treatment (Fig. 3E and 3F). We crossed *pFER:FER-GFP* into *bin2-1*, a dominant gain-of-function allele rendering constitutive repression of BR signaling, and observed decreases in both abundance and plasma membrane distribution of FER-GFP compared to the wild-type background (Fig. 3G-J), whereas qRT-PCR analysis showed no significant difference in *FER* RNA level between *bin2-1* and wild type (fig. S3A). Furthermore, bikinin treatment increased the FER-GFP translocation from ER to PM (fig. S3B-S3D). Taken together, these observations demonstrate that the protein accumulation and ER-to-PM translocation of FER are inhibited by BIN2 and promoted by BR.

**Fig. 3.**
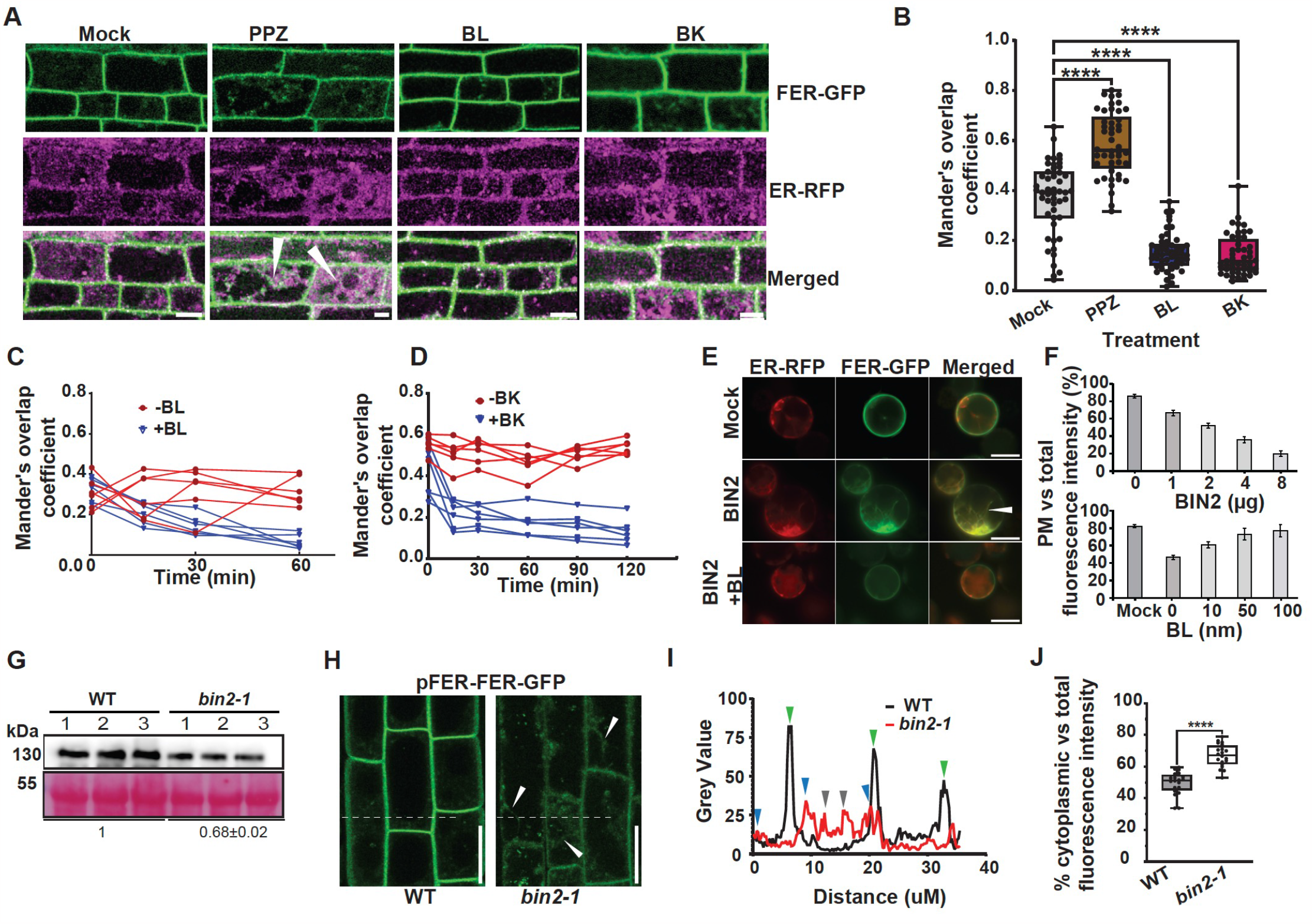
BL promotes and BIN2 inhibits the translocation of FER from ER to PM. (**A**)BR-dependent sub-cellular localization of FER-GFP. The *FER-GFP*/*fer-4*/*ER-RFP* seedlings were grown on solid media under 16 h light/8 h dark cycle for 5 days and then transferred to media supplemented with mock solution, 1 μM propiconazole (PPZ), 10 nM BL, or 30 μM bikinin (BK) for 24 hrs. The epidermal cells of the elongation zone of the root tip were analyzed under a confocal microscope. White arrowheads indicate the colocalization of FER-GFP and ER-RFP. Scale bar = 10 μm. (**B**) Quantification of colocalization FER-GFP and ER-RFP in (A). Values are the mean ± SD of Mander’s overlap coefficient (*33*) of 50 cells from 5 roots. (**C** and **D**) Effects of BL and BK on FER-GFP localization. The seedlings pre-treated with 1 μM PPZ for 24 hours were treated with 1 μM BL (C) and 30 μM BK (D), and the FER-GFP colocalization with ER-RFP was analyzed as in (B) from individual cells (n=6). Four roots were analyzed with similar results. (**E**) *35S:FER-GFP* and *35S:ER-RFP* plasmids were co-transformed with or without *35S:BIN2* in protoplasts and subsequently treated with BL. (**F**) Quantification of (E middle panel) plasma membrane fluorescence as a percentage of total fluorescence intensity after co-transformation with different amounts of BIN2 plasmids (upper panel), or co-transformation with 2 μg *35S:BIN2* plasmids and subsequent BL treatment in protoplasts. Values are the mean ± SD. (**G**) Immunoblot analysis of FER-GFP protein accumulation in *pFER:FER-GFP/fer-4* and *pFER:FER-GFP/fer-4 bin2-1* lines. Quantification is shown as means ± SD of three independent lines. (**H**) Confocal images of *pFER:FER-GFP/fer-4* and *pFER:FER-GFP/fer-4 bin2-1* lines. White arrowheads indicate the ER-like FER-GFP signal in the cytoplasm. Scale bar =20 μm. (**I**) Intensity plot prolife of Image shown in (H) along the white dash line. Green and blue arrowheads indicate the PM signals in WT and *bin2-1*, respectively. Grey arrowheads indicate cytoplasmic FER-GFP signal in *bin2-1*. (**J**) Percentage cytoplasmic versus total fluorescence intensity. Box plot with individual data points of 20 cells per genotype as shown in (H). Three roots were analyzed with similar results. The asterisk indicates a student’s t-test with *p*<0.01.

## BIN2 phosphorylation reduces accumulation and PM localization of FER

Our recent work showed that FER is in proximity to BIN2 and is dephosphorylated upon inhibition of BIN2, identifying FER as a substrate of BIN2 kinase *in vivo* (*25*). To confirm BIN2 phosphorylation of FER, we did *in vitro* phosphorylation of FER cytoplasmic domain (MBP-FER KD) with recombinant BIN2 purified from *E. coli*, followed by mass spectrometry analysis. To avoid FER autophosphorylation, we used a kinase-inactive mutant FER that contains a mutation of the conserved ATP binding pocket (K565R). Mass spectrometry analysis identified phosphopeptides of FER after incubation with wild-type BIN2 but not with kinase-inactive mutant BIN2 (Fig 4A and fig S4A). These include phosphopeptides spanning amino acids 506 to 521 and 871 to 893 in the N-terminal juxta membrane and C-terminal domains, respectively (fig. S4B). Both peptides include the consensus BIN2/GSK3 phosphorylation sites (S/TXXXS/T). The RALF1-induced phosphorylation sites (pS858, pS871, pS874) (*27*) are in proximity to the C-terminal cluster of BIN2 phosphorylation sites (Fig. 4B). The BIN2 phosphorylation sites of FER are evolutionarily conserved in lower plants *Selagenela, Marchantia polymorpha* and *Physcomitrella patens* (Fig. 4B and fig. S4C).

**Fig. 4.**
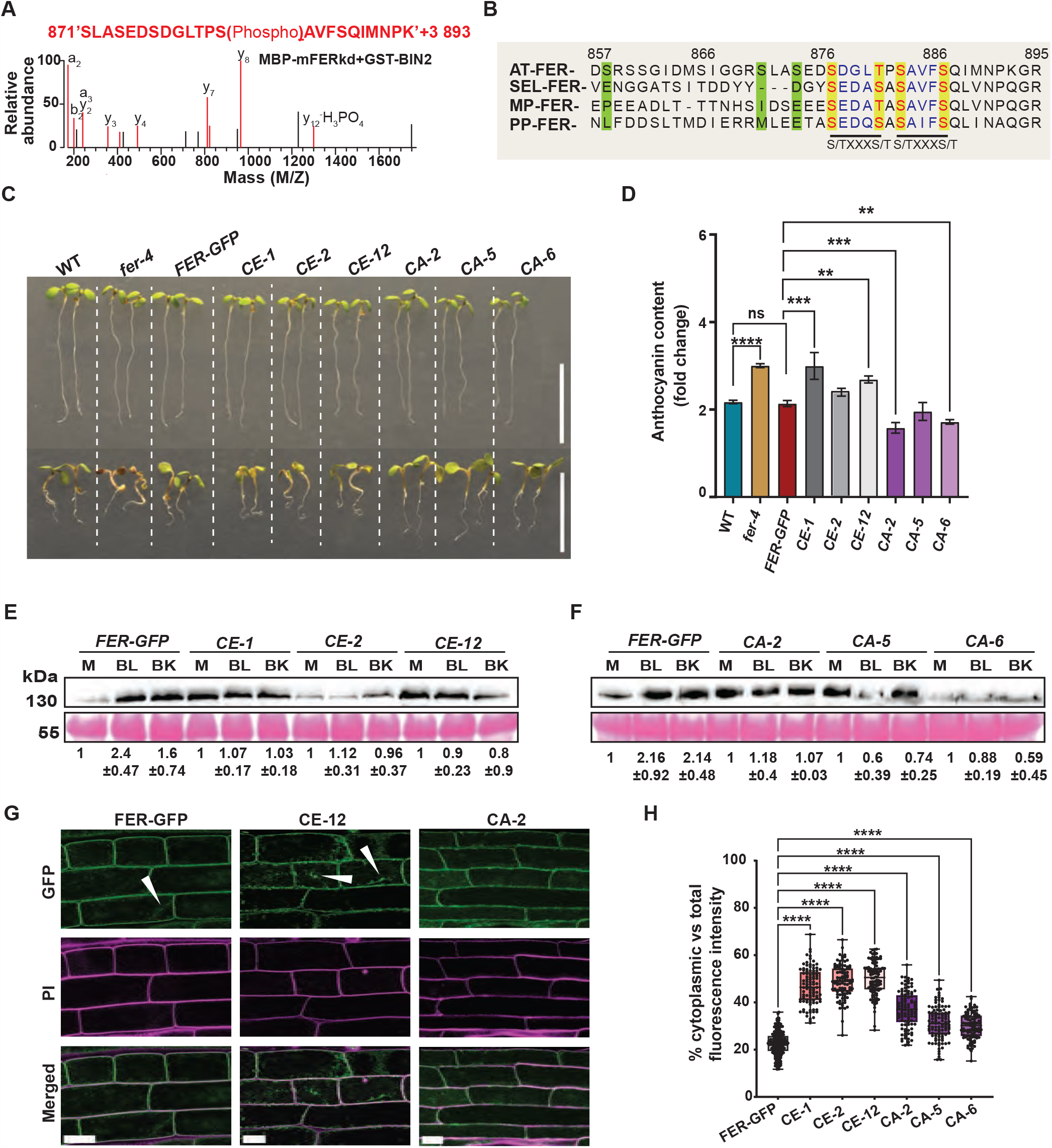
BIN2 phosphorylates FER and inhibits its accumulation and ER-to-PM translocation. (**A**) An example of LC-MS/MS spectra for a FER peptide (aa871-893) phosphorylated by BIN2 *in vitro*. (**B**) Alignment of FER peptide sequences containing BIN2 phosphorylation sites from *Arabidopsis thaliana* (AT-FER), *Selaginella* (SEL-FER), *Marchantia polymorpha* (MP-FER), *Physcomitrella patens* (PP-FER). The conserved BIN2 phosphorylation sites (S/TXXXS/T) are in red text. The RALF1-induced phosphorylation sites are marked by the green boxes. (**C**) Seedlings of WT, *fer-4, pFER:FER-GFP*/*fer-4 (FER_GFP), pFER:FER*^*CE*^-*GFP*/*fer-4 (CE*), and *pFER:FER*^*CA*^*-GFP*/*fer-4* (*CA*) transgenic lines were grown in the absence (upper row) or presence of 1 μM BL (lower row). (**D**) Anthocyanin quantification in samples shown in (C). Values are the means ± SD of three biological replicates. Statistical significance was calculated using one-way ANOVA followed by Tukey’s test. The asterisk indicates the P < 0.0001. (**E** and **F**) Immunoblot analysis of FER-GFP protein in the *pFER:FER-GFP*/*fer-4* (*FER-GFP*), *CE*, and *CA* transgenic lines. Seedlings were grown on solid media for 6 days and then treated for 15 mins with mock solution (M), 1 μM BL, or 30 μM BK. Quantification is shown as means ± SD of three independent lines. (**G**) Subcellular localization of FER-GFP in *pFER:FER-GFP*/*fer-4, CE*, or *CA* plants. Scale bars = 10 μm. (**H**) Quantification of percentage cytoplasmic versus total fluorescence intensity in (G). Box plot with individual data points (WT, N=200 cells/10 roots; CE and CA, N=100 cells/5 roots per line). Statistical significance was calculated using one-way ANOVA followed by Tukey’s test. The asterisks indicate the *P* < 0.0001.

To test the functions of BIN2-mediated phosphorylation of FER, we mutated the two clusters of BIN2 phosphorylation sites at the N-terminal (FER-N: T506, T508, T509, S511, S514, S515, S518) and C-terminal sides (FER-C: S871, S874, 877, T881, S883, S887) of FER kinase domain to either alanine (FER-NA, FER-CA, unphosphorylatable mutations) or to glutamic acid (FER-NE, FER-CE, phosphomimic mutations). We generated transgenic lines that express the wild-type or the mutated FER-GFP in the *fer-4* mutant background. Three independent homozygous lines with comparable transcript and protein levels were selected for each construct (fig. S5, A and B). We observed that FER-CE-GFP lines showed similar BR responses to the *fer-4* mutant, with smaller and reddish seedlings, whereas the FER-CA-GFP plants showed bigger and more greenish seedlings. Quantitation of anthocyanin confirmed that the FER-CE-GFP lines showed elevated anthocyanin levels and FER-CA-GFP phospho-mutants showed reduced anthocyanin levels compared to wild type and the *fer* mutant expressing the wild-type FER (Fig. 4C-D). The plants expressing FER-NE or FER-NA mutations did not show significant differences from the wild type (fig. S5C). These results indicate that phosphorylation of the C-terminal region inhibits FER function.

Finally, we analyzed whether the C-terminal phosphorylation sites mediate BR regulation of the accumulation and PM localization of FER. We analyzed the wild-type and mutant FER-GFP proteins by immunoblotting after treating the transgenic plants with BL or bikinin for 15 min. Contrary to wild-type FER-GFP, the level of FER-CE or FER-CA proteins were not increased by BL treatment (Fig 4E and F), indicating that phosphorylation of these mutated residues mediates BL- and bikinin-enhanced FER-GFP accumulation. Next, we analyzed the subcellular localization of the mutated FER-GFP in the epidermal cells of the elongation and transition zones of the root meristem. We observed that FER-CE-GFP showed increased cytoplasmic GFP-signal compared to wild-type FER-GFP (Fig 4G and H). The FER-CA-GFP showed lower cytoplasmic GFP signals than FER-CE-GFP but slightly higher than the wild type, suggesting that FER localization is dependent on phosphorylation. These results indicate that when the BR level is low, BIN2 phosphorylates the conserved residues in the FER C-terminal region to maintain a low level of FER at the plasma membrane. BR signaling inactivates BIN2, leading to dephosphorylation and PM localization of FER and enhanced surveillance of cell wall damage (Fig. 5).

**Fig. 5.**
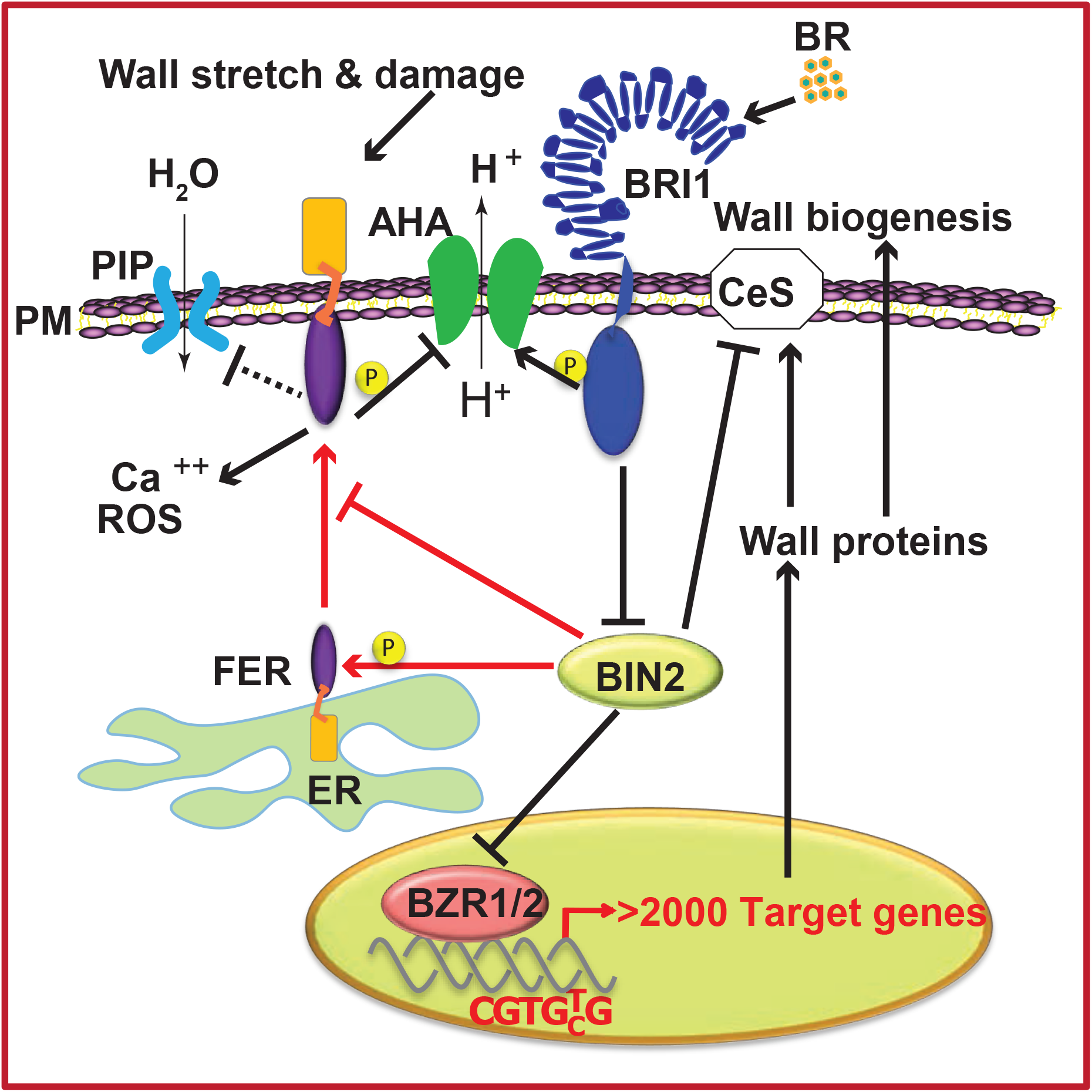
The BR/BIN2-FER signaling circuit ensures integrity during BR-induced cell expansion. BIN2 inhibits cell expansion by phosphorylating and inactivating BZR1 and cellulose synthase (CeS). BIN2 phosphorylates FER to maintain a low level of FER on the PM when the cell is not expanding. BR activates BRI1 leading to BIN2 inactivation, which results in BZR1 accumulation and expression of growth-promoting genes in the nucleus, cellulose synthesis, and accumulation of FER at the PM. BRI1 also phosphorylates the PM H^+^-ATPase to acidify the cell wall and polarize the PM, which loosens the cell wall and increases the turgor pressure, respectively. Upon activation by cell wall damage signals, FER inhibits PM H^+^-ATPase and aquaporin, leading to PM depolarization, cell wall alkalinization, and reduced water uptake, which together rapidly reduce turgor pressure and strengthen cell wall to prevent cell bursting.

## Discussion

Taken together, our study demonstrates a signaling circuit essential for preventing cell rupture during BR-induced cell expansion (Figure 5). When BR level is low, cell elongation is not stimulated and BIN2 phosphorylates FER at its C-terminal conserved phosphorylation sites to maintain a low level of FER at the plasma membrane. When BR is high, BIN2 is inactivated; this not only activates genes and cellular components that drive cell expansion but also increases FER accumulation at the plasma membrane and enhances CWI surveillance. Upon detection of cell wall damage, FER slows down BR-induced cell elongation to prevent further wall damage and cell rupture. In the absence of FER function, BR signaling leads to unabated cell elongation that causes cell wall damage and cell rupture. Mutation of the BIN2 phosphorylation sites in FER abolished BR regulation of FER accumulation and compromised the ability of FER to prevent cell damage during BR-induced growth. Our study demonstrates that the BIN2-FER signaling axis is an essential part of the BR signaling pathway for growth regulation, specifically for maintaining CWI by feedback regulation of the growth rate.

A wide range of phenotypes have been observed in the *fer* mutant. These include defects in pollen guidance and sperm release in sexual reproduction, development of root hair, hormonal responses, mechanic sensing, plant-microbe interactions, and maintenance of CWI under abiotic stresses (*16, 17, 19, 22-24*). These observations have led to the notion that FER participates in or crosstalks with many signaling pathways. In particular, the *fer* mutants show altered responses to many hormones, and FER is considered a central node that mediates crosstalk among multiple phytohormones (*28*). However, our study supports the notion that some of the pleiotropic phenotypes of *fer* may reflect the consequence of losing mechanical sensing and wall integrity rather than a direct role of FER in the specific signaling pathways controlling the phenotypes (*29*).

The mechanisms by which FER slows down BR-induced cell elongation remain to be studied. One possibility is antagonism at the plasma membrane (PM) H^+^-ATPase. BR has been shown to induce activation of PM H^+^-ATPase to reduce cell wall pH and increase turgor pressure, driving rapid cell expansion (*8, 9*). In contrast, FER has been reported to phosphorylate and inhibit PM H+-ATPase and aquaporin (*27, 30, 31*). The antagonism between BR and FER signaling on PM H^+^-ATPase would provide a feedback mechanism for rapid regulation of turgor pressure and cell wall loosening upon detection of wall damage. In addition, FER may inhibit BR signaling, by activating BIN2 or inhibiting BRI1 at the PM, for example, to inhibit BR promotion of cell elongation. Such inhibition of upstream BR signaling, however, would not only have slower effects on turgor pressure and apoplastic pH than the direct FER regulation of PM H^+^-ATPase and aquaporin but also would impede BR-induced processes that enforce the cell wall (*1, 32*). Indeed, cell wall defects have been shown to activate BR signaling as a compensatory mechanism (*10*). We propose that the BR/BIN2-FER signaling circuit fine-tunes the balance between turgor pressure, wall loosening, and wall synthesis to achieve optimal growth while maintaining CWI (Figure 5).

## Acknowledgment

We thank Prof. Kay Schneitz (TU Munich) for the seed of *the1-1, sub21, mik2-1*; Prof. Jose Dinneny (Stanford University) for seed of *fer-4, fer-2, herk1-1, ralf1-1*, pFER-FER-GFP/*fer-4*, and 35S-RALF1 lines. We thank Zhuoran Lyu for technical assistance and May S. Ong for editing the manuscript.

## Author contributions

Conceptualization: A.C., Y.C. H., A.Y.C., and Z.Y.W. Methodology: A.C., Y.C.H., F.L., J.Y. Investigation: A.C., Y.C.H., F.L., and J.Y. Data analysis: A.C., Y.C.H., F.L., J.Y., H.M.W., A.Y.C., S.X., and Z.Y.W. Writing: A.C., A.Y.C., and Z.Y.W. Supervision: H.M.W., A.Y.C., S.X., and Z.Y.W.

## Data and materials availability

All data are available in the main text or supplementary materials. Materials used in the study are available on request.

## Figure Legend

**fig. S1. The *fer* mutant, but not mutants of *FER* homologs, show abnormal responses to BR**. (**A**) Seedlings of WT, *fer-4, ralf1-1, herk1-1, the1-1, sub-21*, and *mik2-1* were grown under 16 h light/8 h dark cycle for 6 days on solid media containing the indicated concentrations of BL. Scale bars are 10 mm. (**B, C**) Quantification of root (B) and hypocotyl (C) lengths of seedlings shown in (A). Results presented here are mean ± SD of measurements pooled from three independent experiments (25≤N≤36 seedlings). Statistical significance was calculated using one-way ANOVA followed by Tukey’s test. Means with different letters are significantly different at P < 0.0001. (**D**) Anthocyanin accumulation in WT, *fer-4, ralf1-1, herk1-1, the1-1, sub-21*, and *mik2-1*. Values are the means ± SD of three biological replicates. The experiment was performed three times with similar results.

**fig. S2. *FER* overexpression attenuates BR-induced growth response.(A)**RT-qPCR analysis of *FER* expression in 6-day-old seedlings of each genotype. Values are the means ± SD of three biological replicates. Statistical significance was calculated using one-way ANOVA followed by Tukey’s test. Asterisk indicates the P < 0.05. (**B**) Seedlings of WT, *fer-4*, three *35S:FER-GFP* (*FER-OE*) lines, *ralf1-1* and *35S:RALF1* (*RALF1-OE*) lines grown on media containing indicated concentrations of BL. Note the greener color of cotyledons and longer roots in *FER-OE* seedlings compared to WT at higher amounts of BL treatment. (**C** and **D**) Quantification of root (B) and hypocotyl (C) length of seedlings shown in (A). Results presented here are mean ± SD of measurements pooled from three independent experiments (25≤N≤37 seedlings). Statistical significance was calculated using one-way ANOVA followed by Tukey’s test. Means with different letters are significantly different at P < 0.0001. Scale bar mm. (**E**) Anthocyanin accumulation in WT, *fer-4, FER-OEs, ralf-1* and *RALF1-OE* lines. Values are the means ± SD of three biological replicates. The experiment was performed three times with similar results.

**fig. S3. The *bin2-1* mutation affects FER-GFP accumulation and sub-cellular localization without affecting the *FER* RNA levels**. (**A**) RT-qPCR analysis of *FER* expression in 6-day-old seedlings of WT, *fer-4*, and heterozygous *bin2-1+/-* lines. Values are the means ± SD of three biological replicates. The asterisk indicates the *p* < 0.05, calculated using one-way ANOVA followed by Tukey’s test. (**B**) Quantification of total GFP fluorescence intensity (AU, artificial unit) of samples shown in Fig. 3H. Box plot with individual data points (19≤N≤20 roots). The asterisk indicates a student’s t-test with *p*<0.01. (**C**) Confocal analysis of FER-GFP in a bin2-1+/-. The epidermal cells from the transition zone of the root tip in 5-day-old *pFER:FER-GFP*/*bin2-1+/-* seedlings were analyzed under a confocal microscope at indicated times before (-BK) and after (+BK) treatment with 50 mM bikinin. The plot profiles show FER-GFP intensity along the white dash line intersecting the nucleus as depicted in the images. The plasma membrane signal is marked with yellow arrowheads (no. 1 and 4) and the cytoplasmic signal is marked by pink arrowheads (no. 2 and 3). Values next to the arrowheads show the FER-GFP fluorescent intensity. Scale bars = 20 μm. (**D**) Quantification of plasma membrane/cytoplasmic fluorescence intensity ratio in (C). Box plot with individual data points (N=15 cells). Statistical significance was calculated using one-way ANOVA followed by Tukey’s test. Means with different letters are statistically significant with each other at *p* < 0.01. The experiment was performed twice with similar results.

**fig. S4. BIN2 phosphorylates FER**. (**A**) An example of LC-MS/MS spectra for a FER peptide (aa506-521) phosphorylated by BIN2 *in vitro*. (**B**) The sequence of FER cytoplasmic domain shows the BIN2 phosphorylation sites detected by LC-MS/MS (red letter) and predicted based on GSK3 consensus target (S/TXXXS/T, dash underlines) in the regions N- and C-terminal to the kinase domain. The previously identified RALF1-induced phosphorylation sites are marked by cyan solid underlines. All S/T residues in the N- and C-terminal regions marked by solid black underlines were replaced with A or E to create the FER^NA^ or FER^NE^ and FER^CA^ or FER^CE^ mutant constructs. (**C**) Sequence alignment of BIN2-site-containing Juxta membrane region N-terminal to the kinase domain of FER from *Arabidopsis thaliana* (AtFER), *Selaginella moellendorffi* (SmFER), *Marchantia polymorpha* (MpFER), and *Physcomitrella patens* (PpFER). The consensus BIN2 sites (S/TXXXS/T) are marked by underlines and the BIN2-phosphorylated S residue detected by LC-MS/MS is marked by red color.

**fig. S5. Transgenic *fer* plants express wild-type and mutant FER-GFP containing mutations of BIN2 phosphorylation sites.(A)**RT-qPCR analysis of *FER* expression in 6-day-old seedlings of *WT, fer-4, pFER:FER-GFP/fer-4 (FER-GFP), pFER:FER*^*CE*^*-GFP*/*fer-4* (*CE*) and *pFER:FER*^*CA*^*-GFP*/*fer-4* (*CA*) transgenic lines. Values are the means ± SD of three biological replicates. Statistical significance was calculated using one-way ANOVA followed by Tukey’s test. Asterisks indicate the P < 0.0001. (**B**) Western blot analysis of FER-GFP protein in *FER-GFP, CE*, and *CA* transgenic lines. (**C**) Seedlings of WT, *fer-4, FER-GFP, pFER:FER*^*NE*^*-GFP*/*fer-4* (*NE*) and *pFER:FER*^*NA*^*-GFP*/*fer-4 (NA*) transgenic lines grown in the absence (-BL) or presence of 1 μM BL (+BL). Scale bar is 10 mm.

## References

1. S. Wolf, Cell Wall Signaling in Plant Development and Defense. Annu Rev Plant Biol 73, 323–353 (2022).

2. J. Chaiwanon, Z. Y. Wang, Spatiotemporal brassinosteroid signaling and antagonism with auxin pattern stem cell dynamics in Arabidopsis roots. Curr Biol 25, 1031–1042 (2015).

3. S. Savaldi-Goldstein, C. Peto, J. Chory, The epidermis both drives and restricts plant shoot growth. Nature 446, 199–202 (2007).

4. R. Kelly-Bellow et al., Brassinosteroid coordinates cell layer interactions in plants via cell wall and tissue mechanics. Science 380, 1275–1281 (2023).

5. J. Y. Zhu et al., The F-box Protein KIB1 Mediates Brassinosteroid-Induced Inactivation and Degradation of GSK3-like Kinases in Arabidopsis. Mol Cell 66, 648–657 e644 (2017).

6. T. W. Kim, Z. Y. Wang, Brassinosteroid Signal Transduction from Receptor Kinases to Transcription Factors. Annu Rev Plant Biol 61, 681–704 (2010).

7. T.-W. Kim et al., Brassinosteroid signal transduction from cell-surface receptor kinases to nuclear transcription factors. Nat Cell Bio 11, 1254–1260 (2009).

8. K. Caesar et al., A fast brassinolide-regulated response pathway in the plasma membrane of Arabidopsis thaliana. Plant J 66, 528–540 (2011).

9. A. Minami, K. Takahashi, S. I. Inoue, Y. Tada, T. Kinoshita, Brassinosteroid Induces Phosphorylation of the Plasma Membrane H+-ATPase during Hypocotyl Elongation in Arabidopsis thaliana. Plant Cell Physiol 60, 935–944 (2019).

10. S. Wolf, J. Mravec, S. Greiner, G. Mouille, H. Hofte, Plant cell wall homeostasis is mediated by brassinosteroid feedback signaling. Curr Biol 22, 1732–1737 (2012).

11. J. M. Escobar-Restrepo et al., The FERONIA receptor-like kinase mediates male-female interactions during pollen tube reception. Science 317, 656–660 (2007).

12. H. Guo et al., Three related receptor-like kinases are required for optimal cell elongation in Arabidopsis thaliana. Proc Natl Acad Sci U S A 106, 7648–7653 (2009).

13. A. Chaudhary et al., Cell wall damage attenuates root hair patterning and tissue morphogenesis mediated by the receptor kinase STRUBBELIG. Development 148, (2021).

14. V. Demidchik et al., Stress-induced electrolyte leakage: the role of K+-permeable channels and involvement in programmed cell death and metabolic adjustment. J Exp Bot 65, 1259–1270 (2014).

15. A. Y. Cheung, Q. Duan, C. Li, M. C. James Liu, H. M. Wu, Pollen-pistil interactions: It takes two to tangle but a molecular cast of many to deliver. Curr Opin Plant Biol 69, 102279 (2022).

16. C. Li, H. M. Wu, A. Y. Cheung, FERONIA and Her Pals: Functions and Mechanisms. Plant Physiol 171, 2379–2392 (2016).

17. W. Feng et al., The FERONIA Receptor Kinase Maintains Cell-Wall Integrity during Salt Stress through Ca(2+) Signaling. Curr Biol 28, 666–675 e665 (2018).

18. Q. Duan, D. Kita, C. Li, A. Y. Cheung, H. M. Wu, FERONIA receptor-like kinase regulates RHO GTPase signaling of root hair development. Proc Natl Acad Sci U S A 107, 17821–17826 (2010).

19. Q. Duan et al., FERONIA controls pectin- and nitric oxide-mediated male-female interaction. Nature 579, 561–566 (2020).

20. C. Liu et al., Pollen PCP-B peptides unlock a stigma peptide-receptor kinase gating mechanism for pollination. Science 372, 171–175 (2021).

21. J. Huang et al., Stigma receptors control intraspecies and interspecies barriers in Brassicaceae. Nature 614, 303–308 (2023).

22. S. Zhong et al., RALF peptide signaling controls the polytubey block in Arabidopsis. Science 375, 290–296 (2022).

23. J. Xing, D. Ji, Z. Duan, T. Chen, X. Luo, Spatiotemporal dynamics of FERONIA reveal alternative endocytic pathways in response to flg22 elicitor stimuli. New Phytol 235, 518–532 (2022).

24. C. Liu, H. Yu, A. Voxeur, X. Rao, R. A. Dixon, FERONIA and wall-associated kinases coordinate defense induced by lignin modification in plant cell walls. Sci Adv 9, eadf7714 (2023).

25. T. W. Kim et al., Mapping the signaling network of BIN2 kinase using TurboID-mediated biotin labeling and phosphoproteomics. Plant Cell 35, 975–993 (2023).

26. C. Li et al., Glycosylphosphatidylinositol-anchored proteins as chaperones and coreceptors for FERONIA receptor kinase signaling in Arabidopsis. Elife 4, (2015).

27. M. Haruta, G. Sabat, K. Stecker, B. B. Minkoff, M. R. Sussman, A peptide hormone and its receptor protein kinase regulate plant cell expansion. Science 343, 408–411 (2014).

28. H. Liao, R. Tang, X. Zhang, S. Luan, F. Yu, FERONIA Receptor Kinase at the Crossroads of Hormone Signaling and Stress Responses. Plant Cell Physiol 58, 1143–1150 (2017).

29. A. Malivert, O. Hamant, Why is FERONIA pleiotropic? Nat Plants, (2023).

30. M. Haruta, W. M. Gray, M. R. Sussman, Regulation of the plasma membrane proton pump (H(+) ATPase) by phosphorylation. Curr Opin Plant Biol 28, 68–75 (2015).

31. J. Bellati et al., Novel Aquaporin Regulatory Mechanisms Revealed by Interactomics. Mol Cell Proteomics 15, 3473–3487 (2016).

32. C. Sanchez-Rodriguez et al., BRASSINOSTEROID INSENSITIVE2 negatively regulates cellulose synthesis in Arabidopsis by phosphorylating cellulose synthase 1. Proc Natl Acad Sci U S A 114, 3533–3538 (2017).

33. K. W. Dunn, M. M. Kamocka, J. H. McDonald, A practical guide to evaluating colocalization in biological microscopy. Am J Physiol Cell Physiol 300, C723–742 (2011).

