## Supplementary material for "Brassinosteroid recruits FERONIA to safeguard cell expansion in Arabidopsis": Chaudhary et a Material and Method.pdf

**Supplementary Materials**

Materials and methods.

Supplementary Figures

### Materials and Methods

#### Plant materials and growth conditions.

Arabidopsis thaliana plants Wild type (WT) used in this study is Columbia ecotype (Col-0) and all mutant lines are in Col-0 background. The *fer-4*, *fer-2*, *FER-GFP/fer-4, ralf1-1*, RALF OE1, (11, 17) , *herk1-1* (12), *the1-1*, *sub-21*, *mik2-1* (13) *bin2-1* (5) lines were previously described. FER-GFP/*fer-4 bin2-1* were generated by crossing FER-GFP/*fer-4* into *bin2-1.* Seedlings were grown on 0.9% phytoblend agar medium containing ½ Murashige and Skoog basal medium (MS) (pH5.7) and 1% sucrose under 16-hour -ight/8-hour-dark cycle in a growth chamber (Percival) at 22$^{o}$C.

#### BL and bikinin treatment and seedling growth assays

Seeds were sterilized by 75% (V/V) ethanol and stratified at 4$^{o}$C for 48 hours. Then grown on solid 1/2 MS medium (as described above) supplemented with DMSO mock or Brassinolide (BL) or Bikinin on a 16/8-hour light/dark cycle at 22$^{0}$C in the growth chamber for 6 days. The hypocotyl, and root length was measured by ImageJ. For growth rate analysis, Seedlings were first grown on 1/2 MS with 1% sucrose 0.9% agar plate containing 1 μM propiconazole (PPZ) for 4 days, then transferred to MS1/2 with 1% sucrose 0.9% agar plate with either mock (DMSO plus 1 μM PPZ) or 1 μM BL plus 1 μM PPZ. High-resolution time-lapse images were acquired with an in-house vertical imaging system.

#### Anthocyanin extraction and quantification

BL- or mock-treated seedlings were harvested in liquid nitrogen and grounded in tissue homogenizer. The homogenized tissue was suspended in five volumes (based on fresh weight) of 45% methanol and 5% acetic acid. After a short vortex, samples were stored at 4$^{0}$C for 12 hours in the dark then centrifuged at 14,000 rpm for 5 mins. The clear pink color supernatant was collected. The relative level of anthocyanin was calculated from the absorbance at 530 and 637 nm (Abs 530-0.25×Abs 657)/fresh weight).

#### Electrolyte leakage assay.

For electrolyte leakage, seedlings were first grown on 1/2 MS with 1% sucrose 0.9% agar plate containing 1 μM propiconazole (PPZ) for 4 days, then 20 seedlings from each genotype were transferred to deionized water (dd$H_2O$) plus 1% sucrose with 1 μM PPZ for 3 hours to get habituated. The medium was exchanged from habituated seedlings with fresh $ddH_2O$ and either mock (DMSO plus 1 μM PPZ) or 1 μM BL plus 1 μM PPZ and the conductivity of solution was measured. To measure the total electrolyte of the cell, seedlings were boiled at $954^0$C for 5 mins, briefly centrifuged and conductivity was recorded.

**Site-directed mutagenesis.**

The FER-GFP plasmid the pCAMBIA vector [17] was used as a template for site-directed mutagenesis following the manufacturer's protocol (New England Biolabs) to generate *FER-GFP–S-A/E*, which transformed in *fer-4* mutant plants.

**Transgenic plants**

The *FER-GFP–S-A/E* plasmid was transformed into *fer-4* mutants plants using Agrobacterium mediated floral dipping method. Transformants T1 were selected on Kanamycin antibiotics and further propagated for downstream analysis. For the *FER-GFP/fer-4 ER-RFP* line, transgenic plants *FER-GFP/fer-4* were retransformed with ER-RFP plasmid and selected for double transgene. *FER-GFP/fer-4bin2-1* plants were generated by crossing *FER-GFP/fer-4* into *bin2-1* and F2 lines showing FER-GFP expression and *bin2-1* phenotype were selected.

**Real-time quantitative PCR and Primers**

Total RNA was extracted from fresh seedlings using the Spectrum Plant Total RNA kit (Sigma-STRN250). To prepare cDNA, M-MLV reverse transcriptase (Fermentas) was used and performed as recommended. The quantitative real-time PCR was done using LightCycler 480 (Roche) and the SYBR Green Master Mix (Bioline) following user's guidelines. *Ubiquitin 10* (*UBQ10*) was used as an internal reference. The Primers used in qPCR are provided in supplementary table 1.

**Sequence alignment**

The multiple sequence alignment analysis was performed with Geneious (Geneious Prime® 2022.2.2).

**Protein extraction and immunoblot analysis.**

For protein extraction, plants were frozen in liquid nitrogen, ground, weighed, and added into corresponding 2× SDS buffer (0.125 mM Tris-HCl [pH 6.8], 4% SDS, 20% Glycerol and 2% β-mercapto-ethanol). Samples were heated for 15 min at 65°C, centrifuged at 10000g for 10 min, separated on a 10% (detecting BZR1 protein) polyacrylamide gel and then blotted on Nitrocellulose membranes (Bio-Rad 0.45uM) in 192 mM glycine and 25 mM Tris-HCl with a Trans-blot Turbo blotting system (Bio-Rad) for 15 min. Membranes were blocked for 24 hour at $4^0$C in a blotting buffer (140 mM NaCl, 10 mM KCl, 8 mM Na2HPO4, 2 mM KH2PO4, 0.5% skim milk, and 0.1% Tween20, pH 7.4). The gel blots were incubated with the primary antibodies (anti-GFP, Transgene, HT801) at 1:2000. The secondary antibodies were used at 1:5000 dilutions for 1 hour.

***In vitro* phosphorylation assay and mass spectrometry analysis**

The *in vitro* phosphorylation assay using recombinant proteins was performed following the published procedure [5, 7]. One microgram of GST-BIN2, GST-mBIN2 was incubated with MBP-FER-KD, MBP-mFER-KD, MBP in phosphorylation buffer (20mM Tris pH7.5, 1mM MgCl2, 100 mM NaCl and 1 mM DTT) containing 200 μM ATP at 30 °C for 3 h. The reaction was stopped by heat inactivation at 95 $^{0}$C for 5 min, 2× SDS sample buffer was added and separated by SDS–PAGE, followed by colloidal blue staining (Invitrogen). Desired proteins were excised and subjected for in-gel digestion with trypsin. Peptide mixtures were desalted using C18 ZipTips (Millipore). Data were acquired in the Data Dependent Acquisition (DDA) mode. Briefly, peptides were analysed by liquid chromatography–tandem MS (LC–MS/MS) on a Nanoacquity ultraperformance liquid chromatography system (Waters) connected to a Linear Trap Quadrupole Orbitrap Velos mass spectrometer (Thermo). Peptides were separated using analytical Easy-Spray C18 columns (75 μm × 150 mm) (Thermo, ES800). The flow rate was 300 nl min$^{-1}$, and peptides were eluted by a gradient from 2% to 30% solvent B (acetonitrile/0.1% formic acid) over 57 min, followed by a short wash at 50% solvent B. After a precursor scan was measured in the Orbitrap by scanning from mass-to-charge ratio 350 to 1,400 at a resolution of 60,000, the six most intense multiply charged precursors were selected for collision-induced dissociation in the linear ion trap.The MS/MS data were converted to peaklist using a script PAVA (a peaklist generator that provides a centroid MS2 peaklist), and the data were searched using Protein Prospector against the user protein. A precursor mass tolerance was set to 20 ppm, and MS/MS2 tolerance was set to 0.6 Da. Carbamidomethylcysteine was searched as a constant modification. Variable modifications include protein N-terminal acetylation, peptide N-terminal Gln conversion to pyroglutamate and Met oxidation. Subsequent searches were performed to find those peptides modified by phosphorylation. The search parameters were as above, but this time allowing for phosphorylation on serine, threonine, and tyrosine.

**Microscopy**

For scanning electron microscopy, seedlings were fixed in 100% (V/V) methanol for 30 mins and then washed with 100% (V/V) ethanol for 3 times. The critical point drying (CPD) and gold sputtering was performed as described [13]. The images were acquired with FEI Quanta 200

FEG fitted with OXFORD instruments 6650 EDS detector. For Confocal laser scanning microscopy, a Leica TCS SP8 X microscope equipped with GaAsP (HyD) detectors was used. The objective lenses used for scanning were a water-corrected ×63 objective (NA 1.2), a ×40 objective (NA 1.1), and a ×20 immersion objective (NA 0.75). The scan speed was at 400 Hz, with 4-line average. GFP fluorescence excitation was performed at 488 nm using a multi-line argon laser (3% intensity) and detected at 502-536 nm. RGP fluorescence was excited using a 561 nm laser (1% intensity) and detected at 610-672 nm. For the direct comparisons of fluorescence intensities, laser, pinhole, and gain settings of the confocal microscope were kept identical when capturing the images from the seedlings of different treatments or genotype. For determination of colocalization, Mander's overlap coefficient was calculated using ImageJ/Fiji software (33). Images were adjusted for color and contrast using ImageJ/Fiji software.

**Statistics**

The statistical analysis was performed with Graph pad Prism (PRISM9) software. The details of analysis are described in respective figure legends.
