## Supplementary material for "Brassinosteroid recruits FERONIA to safeguard cell expansion in Arabidopsis": Supplementary figures.pdf

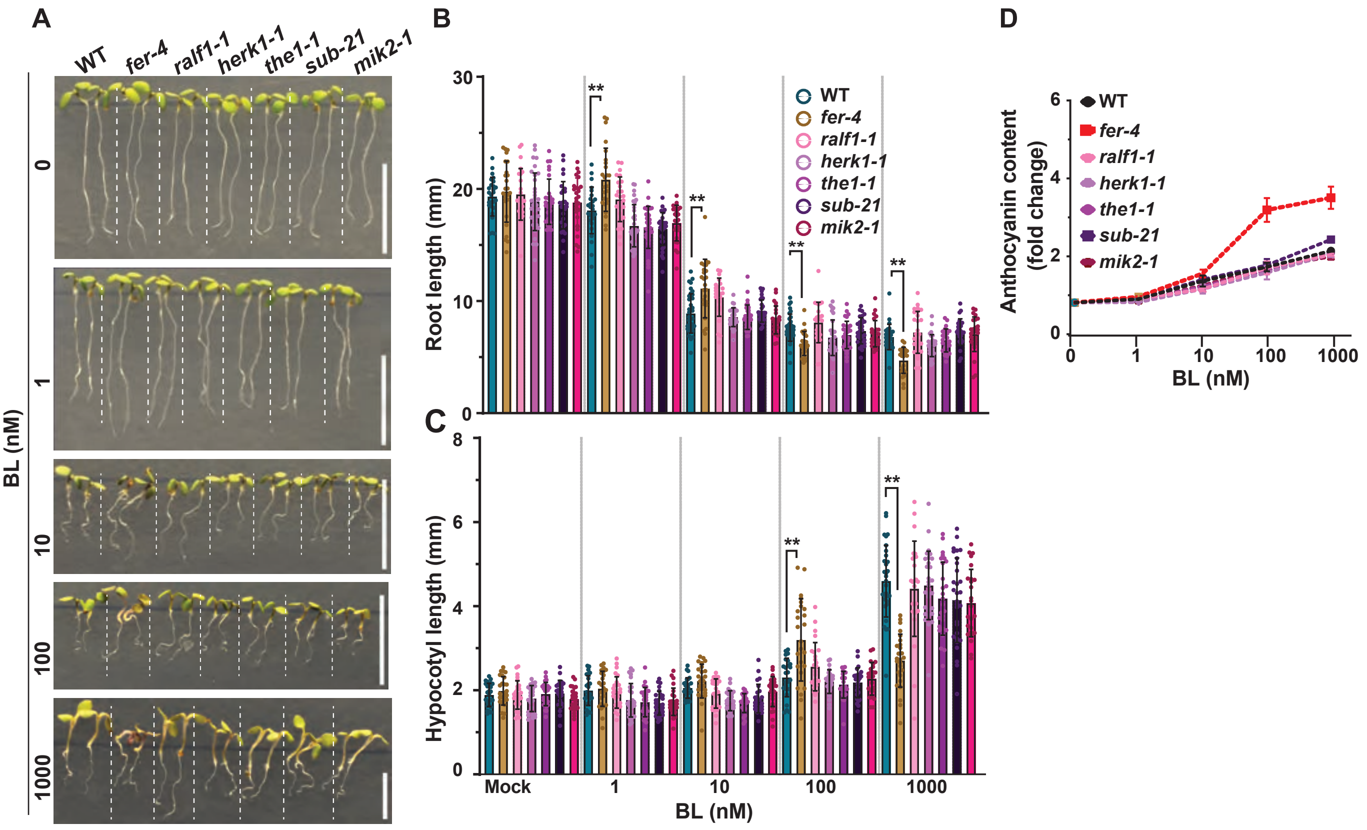


**fig. S1. The *fer* mutant, but not mutants of FER homologs, show abnormal responses to BR.** (**A**) Seedlings of WT, *fer-4, ralf1-1, herk1-1, the1-1, sub-21*, and *mik2-1* were grown under 16 h light/8 h dark cycle for 6 days on solid media containing the indicated concentrations of BL. Scale bars are 10 mm. (**B, C**) Quantification of root (B) and hypocotyl (C) lengths of seedlings shown in (A). Results presented here are mean ± SD of measurements pooled from three independent experiments (25≤N≤36 seedlings). Statistical significance was calculated using one-way ANOVA followed by Tukey's test. Means with different letters are significantly different at P < 0.0001. (**D**) Anthocyanin accumulation in WT, *fer-4, ralf1-1, herk1-1, the1-1, sub-21,* and *mik2-1*. Values are the means ± SD of three biological replicates. The experiment was performed three times with similar results.

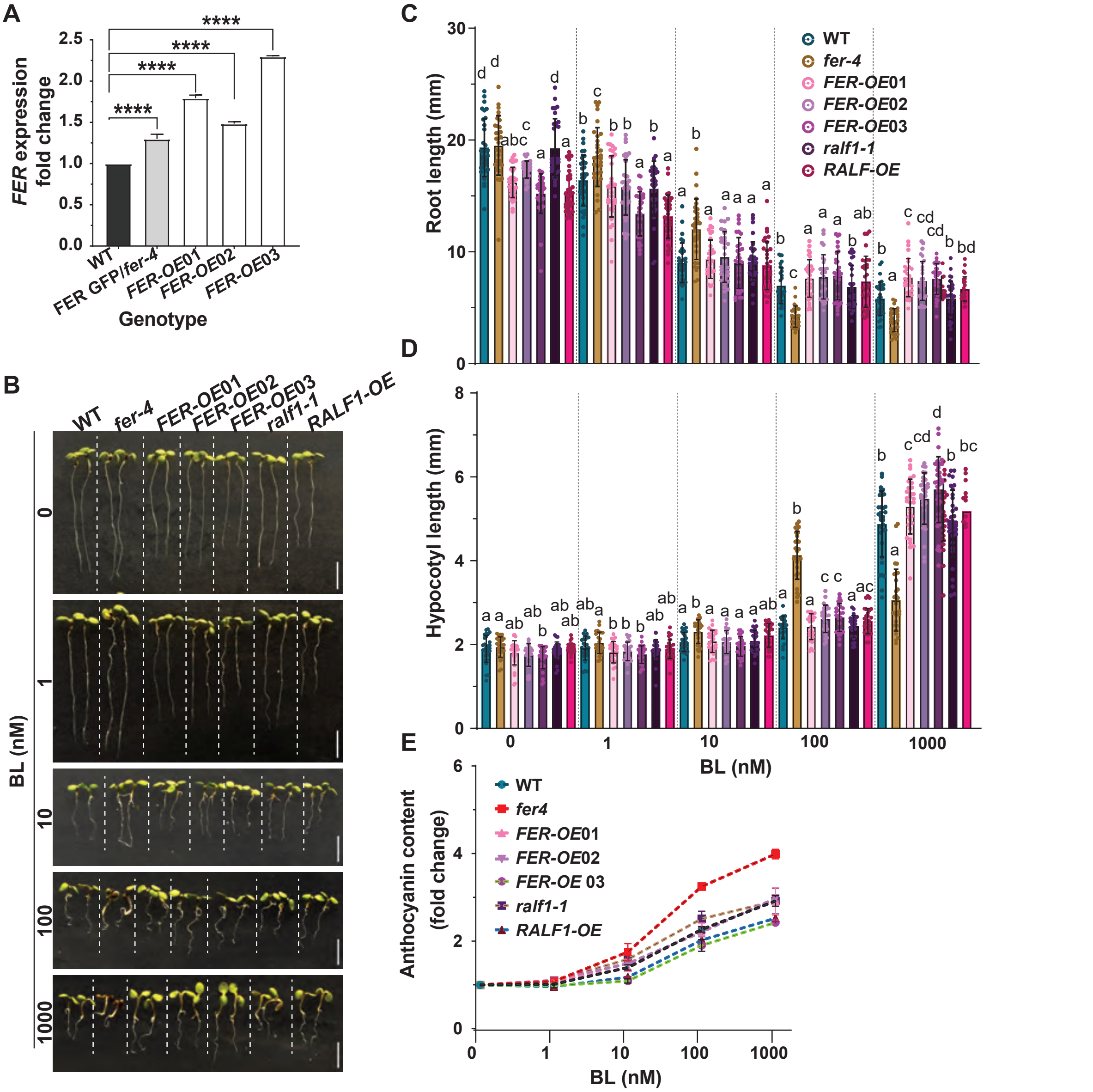


**fig. S2. FER overexpression attenuates BR-induced growth response.**
(**A**) RT-qPCR analysis of FER expression in 6-day-old seedlings of each genotype. Values are the means ± SD of three biological replicates. Statistical significance was calculated using one-way ANOVA followed by Tukey's test. Asterisk indicates the P < 0.0001. (**B**) Seedlings of WT, fer-4, three *35S:FER-GFP (FER-OE)* lines, *ralf1-1* and *35S:RALF1* (*RALF1-OE*) lines grown on media containing indicated concentrations of BL. Note the greener color of cotyledons and longer roots in *FER-OE* seedlings compared to WT at higher amounts of BL treatment.
(**C** and **D**) Quantification of root (B) and hypocotyl (C) length of seedlings shown in (A). Results presented here are mean ± SD of measurements pooled from three independent experiments (25≤N≤37 seedlings). Statistical significance was calculated using one-way ANOVA followed by Tukey's test. Means with different letters are significantly different at P < 0.0001. Scale bar 10 mm. (**E**) Anthocyanin accumulation in WT, *fer-4, FER-OE, ralf-1* and *RALF1-OE* lines. Values are the means ± SD of three biological replicates. The experiment was performed three times with similar results.

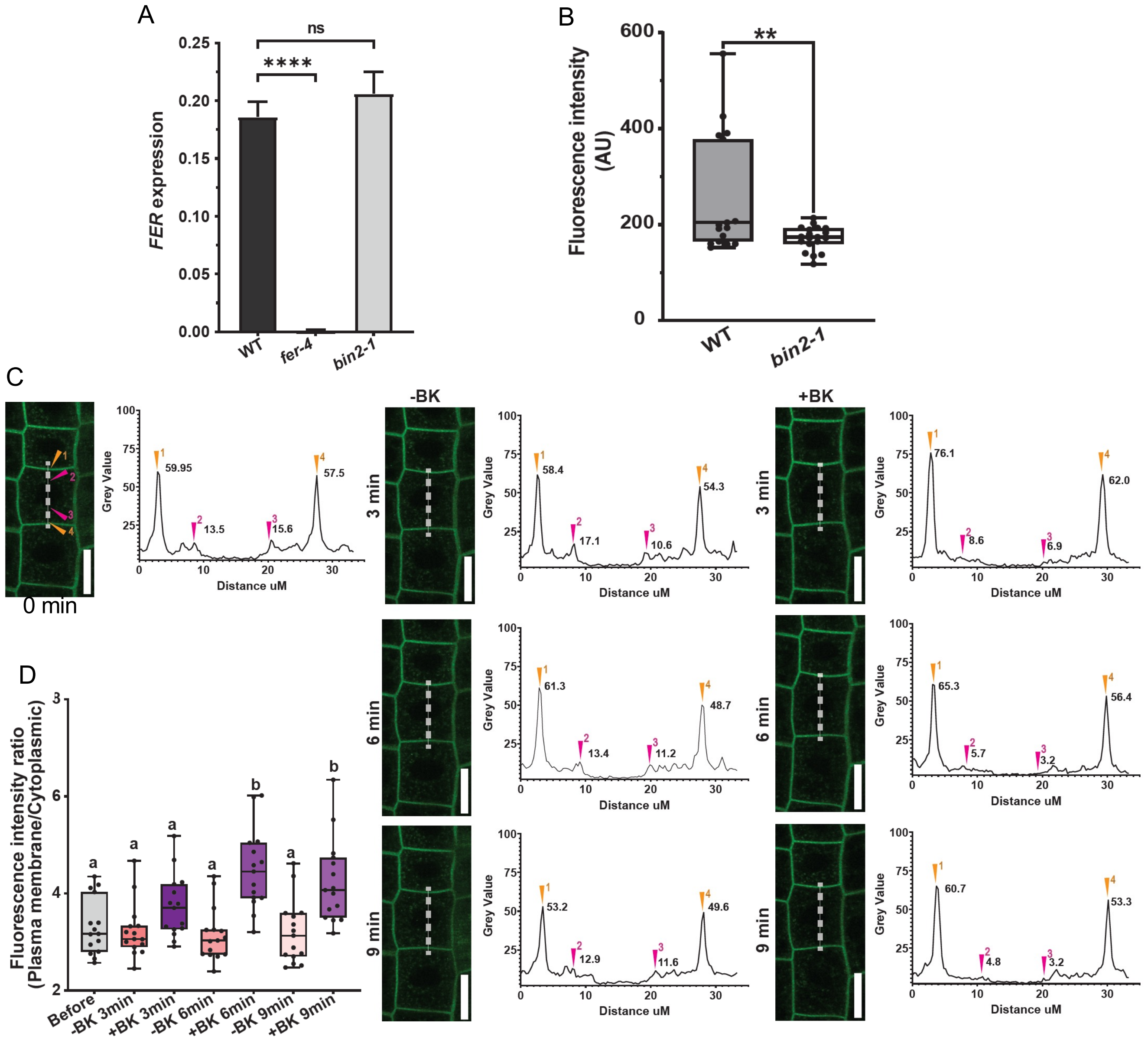


**fig. S3. The *bin2-1* mutation affects FER-GFP accumulation and sub-cellular localization without affecting the *FER* RNA levels.** (**A**) RT-qPCR analysis of *FER* expression in 6-day-old seedlings of WT, *fer-4,* and heterozygous *bin2-1+/-* lines. Values are the means ± SD of three biological replicates. The asterisk indicates the $p<0.05$, calculated using one-way ANOVA followed by Tukey's test. (**B**) Quantification of total GFP fluorescence intensity (AU, artificial unit) of samples shown in Fig. 3H. Box plot with individual data points (19≤N≤20 roots). The asterisk indicates a student's t-test with $p<0.01$. (**C**) Confocal analysis of FER-GFP in *bin2-1+/-.* The epidermal cells from the transition zone of the root tip in a 5-day-old *pFER:FER-GFP*/*bin2-1+/-* seedlings were analyzed under a confocal microscope at indicated times before (-BK) and after (+BK) treatment with 50 μM bikinin. The plot profiles show FER-GFP intensity along the white dash line intersecting the nucleus as depicted in the images. The plasma membrane signal is marked with yellow arrowheads (no. 1 and 4) and the cytoplasmic signal is marked with pink arrowheads (no. 2 and 3). Values next to the arrowheads show the FER-GFP fluorescent intensity. Scale bars = 20 μm. (**D**) Quantification of plasma membrane/cytoplasmic fluorescence intensity ratio in (C). Box plot with individual data points (N=15 cells). Statistical significance was calculated using one-way ANOVA followed by Tukey's test. Means with different letters are statistically significant with each other at $p < 0.01$. The experiment was performed twice with similar results.

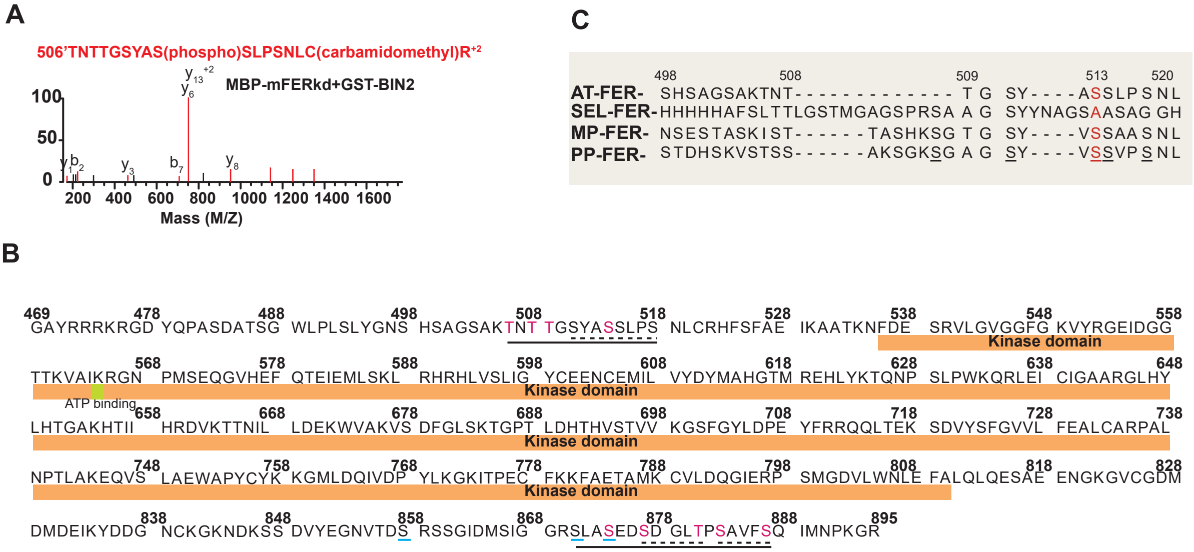


**fig. S4. BIN2 phosphorylates FER.**
(**A**) An example of LC-MS/MS spectra for a FER peptide (aa506-521) phosphorylated by BIN2 *in vitro*. (B) Analysis of BIN2 phosphorylation sites (S/TXXXS/T) in FER cytoplasmic domain. The regions containing BIN2/GSK3 phosphorylation sites are marked by black dash underlines and the previously identified RALF1-induced phosphorylation sites are marked by cyan solid underlines. All S/T residues in the N- and C-terminal regions marked by solid black underlines were replaced with A or E to created the FER$^{NA}$ or FER$^{NE}$ and FER$^{CA}$ or FER$^{CE}$ mutant constructs. (C) Sequence alignment of N-terminal region of FER cytoplasmic domain from Arabidopsis thaliana (AT-FER), Selaginella (SEL-FER), Marchantia polymorpha (MP-FER), Physcomitrella patens (PP-FER). BIN2 target phosphosites (S/TXXXS/T) are in red letters.

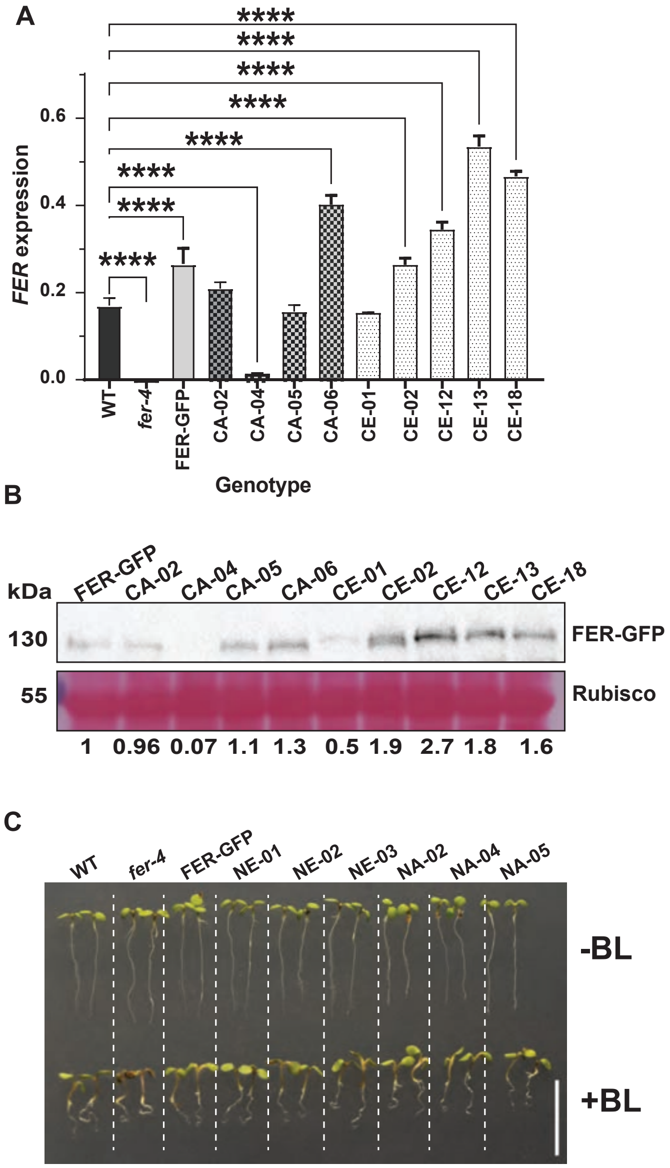


**fig. S5. Transgenic *fer* plants express wild-type and mutant FER-GFP containing mutations of BIN2 phosphorylation sites.**

(**A**) RT-qPCR analysis of *FER* expression in 6-day-old seedlings of WT, *fer-4*, *pFER:FER-GFP/fer-4* (*FER-GFP*), *pFER:FER*$^{CE}$*-GFP/fer-4* (*CE*) and *pFER:FER*$^{CA}$*-GFP/fer-4* (*CA*) transgenic lines. Values are the means ± SD of three biological replicates. Statistical significance was calculated using one-way ANOVA followed by Tukey's test. Asterisks indicate the $P < 0.0001$. (**B**) Western blot analysis of FER-GFP protein in *FER-FER*, *CE*, and *CA* transgenic lines. (**C**) Seedlings of WT, *fer-4*, *pFER:FER-GFP/fer-4*, *pFER:FER*$^{NE}$*-GFP/fer-4* (*NE*) and *pFER:FER*$^{NA}$*-GFP/fer-4* (*NA*) transgenic lines grown in the absence (-BL) or presence 1 μM BL (+BL). Scale bar is 10 mm.
